# Conserved Function of Drg GTPases in Promoting Protein Synthesis in Stalled Ribosomes

**DOI:** 10.1101/2024.04.20.590341

**Authors:** Christopher W. Hawk, Hong Jin

## Abstract

Maintaining proper protein homeostasis is essential for cell physiology. The ribosome and GTPases, which are two of the most ancient and critical cellular molecules, are central players in protein synthesis and its regulation. Here we report the discovery of a new general translation factor that targets stalled ribosomes and promotes protein synthesis in an evolutionarily conserved manner. We show that the essential bacterial Obg GTPases are distant homologs of eukaryotic and archaeal Drg proteins and serve critical roles in promoting efficient protein translation in stalled ribosomes. Through *in vivo* characterization, including cross-species complementation of cells where ribosomes are induced to stall by addition of either the antibiotic anisomycin or exogenous mRNA harboring a long poly(A) sequence, we demonstrate that a conserved function of Drg proteins is to alleviate ribosomal stalling during translation. Our data show that bacterial Obg rescues stalled ribosomes in both Saccharomyces cerevisiae and human cells lacking endogenous Drgs, as does supplementation of the respective endogenous Drg proteins from yeast and human cells. Furthermore, the presence of ObgE and GTP stimulates peptidyl transfer, the key catalytic function of the ribosome, suggesting a possible molecular mechanism of this GTPase to enhance translation in stalled ribosomes. This discovery shows that the Drg protein is a new general translation factor that directly affords cells from the three domains of life a new form of translation regulation.

## Introduction

Proteins play critical functions in nearly all biological processes, and all cellular proteins are synthesized in the ribosome using the nucleotide sequence of messenger RNAs (mRNAs) as a template. Protein synthesis is well appreciated to be tightly regulated to meet the intrinsic and ever-changing demands of cell physiology and function. As a multi-stage process, protein synthesis is controlled at each phase, including translation initiation, elongation, and termination (extensively discussed in many reviews such as (D’Orazio and Green, 2021; Dever et al., 2023; Jackson et al., 2010; Richter and Coller, 2015)).

Decades of research have elegantly uncovered many molecular details of the translation process, including the intrinsic requirement that ribosomes briefly pause at each codon on the mRNA to carry out template-dependent translation. Beyond these natural periodic pauses, ribosomes do not translate the mRNA at a uniform speed (Ingolia et al., 2009; Pedersen, 1984). They slow down, temporarily halt, and may permanently stall over the course of translation. Translation of certain amino acids (e.g. proline, arginine, lysine) (Charneski and Hurst, 2013; Schuller et al., 2017; Zeng et al., 2021), limited availability of amino acids and charged tRNAs (Plotkin and Kudla, 2011; Schuller and Green, 2018), stalling sequences of the nascent peptides in the exit tunnel of the ribosome (Ramu et al., 2011; Woolstenhulme et al., 2013), secondary structures in the mRNA (Doma and Parker, 2006), and interactions with RNA-binding proteins (Darnell et al., 2011) are all demonstrated to modulate the speed of translation.

Despite the potential for attenuated protein output, a subset of translation pauses are programmed and are functionally important. Rare codons are encoded in mRNA sequences to facilitate co-translational protein folding and modifications (Pechmann and Frydman, 2013; Stein and Frydman, 2019; Yu et al., 2015) and sub-cellular localization (Pechmann et al., 2014; Voorhees et al., 2014). Translation pauses may even allow cells to integrate multiple signals and coordinate different pathways in response to internal cues or stress (Jin lab, manuscript in preparation).

On the other end of the spectrum, a prolonged pause of the ribosome on an mRNA results in premature arrest of protein synthesis in the open reading frame, which is conceivably deleterious to the cell if left unresolved because it could generate truncated proteins and also diminish the cellular availability of translating ribosomes. Therefore, a premature translation arrest elicits quality control mechanisms that recycle stalled ribosomes and degrade the incomplete nascent polypeptide chain as well as troubled mRNAs (Brandman and Hegde, 2016; Doma and Parker, 2006; Inada and Beckmann, 2024; Joazeiro, 2019).

Given that ribosomes move unidirectionally from the 5’ to 3’-end of a given mRNA, and mRNAs are frequently translated by multiple ribosomes simultaneously, extended ribosome pauses often lead to collision of two or more translating ribosomes. Notably, cells undergoing fast proliferation and growth often produce an increased number of mature ribosomes. The elevated cellular ribosome concentration leads to increased ribosome flux on a given mRNA, undoubtedly increasing the likelihood of ribosome collisions. It was estimated that around 6-10% of translating ribosomes are in a collided state (Arpat et al., 2020). Given the sheer number of these events, it seems too costly for the cell to commit to quality control on every ribosome that is paused or collided with during translation.

A stochastic ribosome pause or collision, as an inherent feature of the translational process, is likely to be resolved naturally due to its transient nature. However, how does a cell distinguish between programmed and problematic ribosome pauses or collisions? A programmed ribosome pause or collision, such as pauses that are beneficial to protein folding and localization, requires translation machinery to recover and return to the translation pool. On the other hand, a problematic ribosome pause and collision caused by faulty translational components requires activation of quality control mechanisms, such as ribosome-associated quality control (RQC), to disassemble the troubled translation machinery and to degrade the truncated protein (Brandman and Hegde, 2016; Joazeiro, 2019; Sitron and Brandman, 2020). Furthermore, as ribosome pauses and collisions are nearly inevitable, how does the cell prevent premature activation of quality control pathways on a programmed ribosome pause or collision? These are important questions to be answered in the field of molecular biology.

In this study, we report the evolutionarily conserved function of <u>d</u>evelopmentally regulated <u>G</u>TP-binding (Drg) proteins in promoting protein synthesis of stalled ribosomes. Drg proteins are an ancient family of important but poorly understood GTPases that are highly expressed in actively growing and developing cells of plants, animals, and humans (Westrip et al., 2021). The Drg family of proteins was named as such for a historical reason: it was originally characterized as a developmentally regulated protein, as Drg mRNA was highly expressed in neural precursor cells in the developing mouse brain (Kumar et al., 1992). Shortly after its discovery, Drg mRNAs and proteins were found to be widely expressed at variable levels in cultured cell lines, as well as embryonic, postnatal, and adult tissues (Sazuka et al., 1992). An earlier phylogenetic study revealed that eukaryotes typically contain two Drg genes, Drg1 and Drg2, whereas archaea only contain one (Li and Trueb, 2000), which is presumably an ancestral homolog of their eukaryotic counterparts. The amino acid sequences of eukaryotic Drg1 and Drg2 are highly homologous to each other. Both Drg1 and Drg2 proteins contain a canonical G domain, which is comprised of the five conserved G motifs (G1-G5) common to the GTPase Obg-Hflx superfamily (Leipe et al., 2002).

It was reported that Drg proteins directly bind to partner proteins, Dfrp proteins (<u>D</u>rg <u>f</u>amily regulatory <u>p</u>rotein), in eukaryotic cells (Ishikawa et al., 2009; Ishikawa et al., 2005). Drg and Dfrp association confers stability to the Drg protein *in vivo* (Ishikawa et al., 2005) and enhances the GTPase activity of the Drg *in vitro* (Francis et al., 2012; O’Connell et al., 2009). Compared to those of Drg proteins, amino acid sequences of the Dfrp proteins are less conserved and show more inter-species sequence divergence. We recently discovered that yeast Drg complexes suppress ribosome pauses and promote protein synthesis in stalled ribosomes (Zeng et al., 2021), demonstrating that this family of proteins functions at the crossroads of protein synthesis and quality control in eukaryotic cells. Here we extend the conservation of Drg from the eukaryotic and archaeal to the bacterial domain. We report that the bacterial Drg ortholog, Obg, an essential protein in bacteria (Kint et al., 2014; Leipe et al., 2002), rescues the growth and translation defects in yeast and human cells that are caused by deletion of their endogenous Drg proteins. Further, our data show that Obg facilitates puromycin incorporation in stalled ribosomes. Taken together, these results demonstrate that Drg proteins are new conserved general translation factors that function to promote protein synthesis when ribosomes stall from bacteria to humans.

## Results

### Cellular functions of Drg proteins are conserved from bacteria to eukaryotes

Our phylogenetic study shows that *<u>Drg GTPases are conserved across all three domains</u> <u>of life</u>*. The proteins have co-evolved with the ribosome, as any organism or sub-cellular organelle that contains its own ribosomes also contains Drg proteins (unpublished data). The bacterial homolog of Drg, Obg, plays critical functions in DNA replication, chromosome segregation, ribosome assembly and cellular stress response pathways (Kint et al., 2014; Leipe et al., 2002). Although Drg proteins are known to bind to their partner Dfrp in eukaryotic cells, we have not yet uncovered a Dfrp-like protein that is conserved in bacteria using phylogenetic tools.

To determine whether the cellular function of Drg proteins is conserved across the three domains of life, we studied cell growth behavior in the presence of the bacterially derived antibiotic anisomycin, which stalls protein biosynthesis by inhibiting peptidyl transferase activity of the ribosome (Grollman, 1967; Hummel and Bock, 1987). *S. cerevisiae* cells lacking their native Drg1 (Rbg1) are observably less viable in the presence of anisomycin when compared to WT cells (Zeng et al., 2021). Here, we expressed yeast (*S. cerevisiae*), bacterial (*E.coli*), archaeal (*Methanosarcina acetivorans),* and human orthologs of Drg1, namely Rbg1, ObgE, aDrg, and hDrg1, on a constitutively active promoter in yeast cells lacking the endogenous Rbg1 gene and assayed their growth in the presence of anisomycin.

As expected, when compared to WT *S. cerevisiae*, *Δrbg1 S. cerevisiae* exhibited significantly reduced cellular viability when grown in the presence of 12.5µg/mL anisomycin (**Fig 1**. Line 1 and 2). *Δrbg1* yeast expressing vector encoded WT Rbg1 restores the cell growth (**Fig 1**. Line 3).

**Fig. 1.**
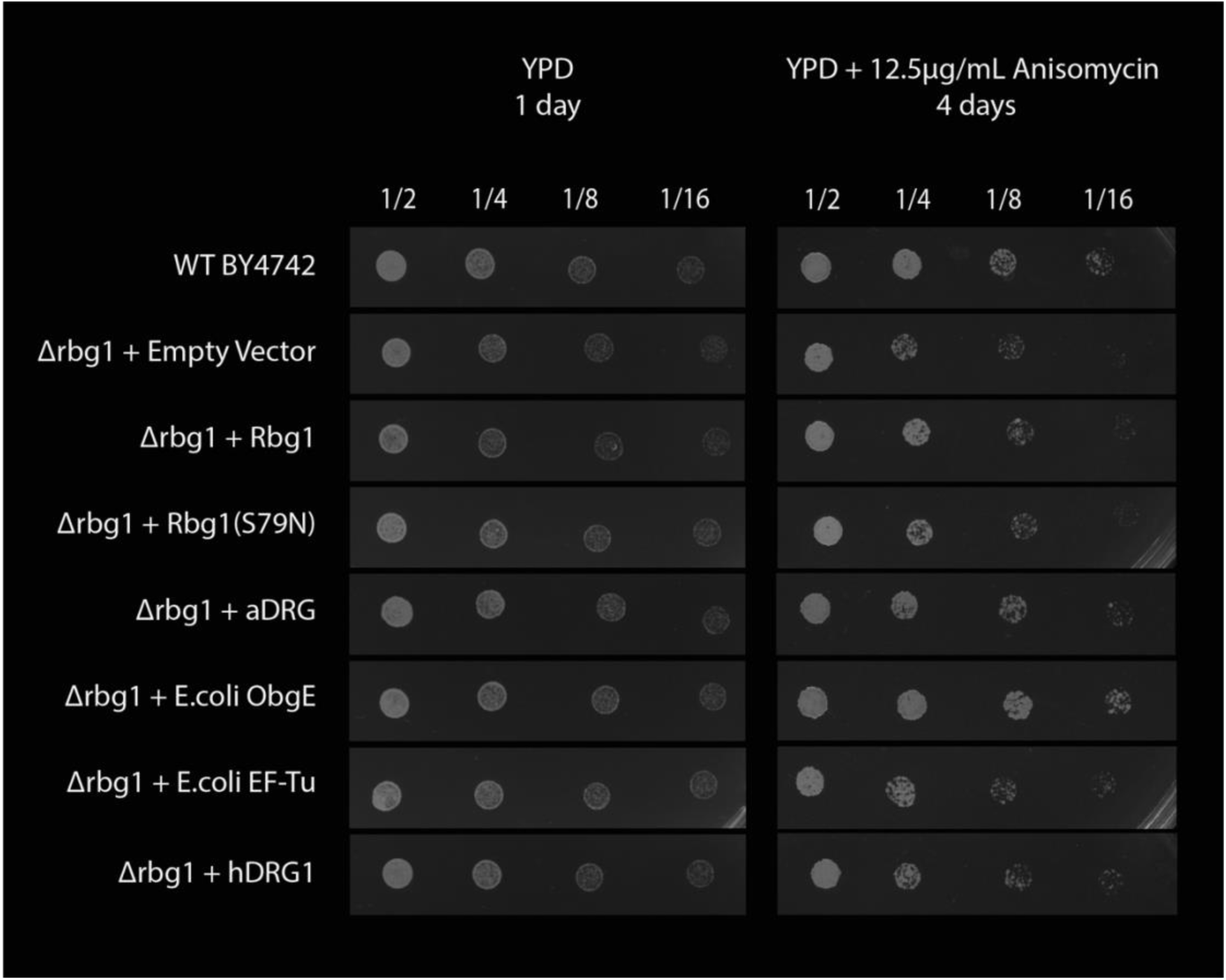
Bacterial and Archaeal Drg proteins compensate for the lost Drg function in the yeast cell. Serial dilutions of wild type and *Δrbg1* S. cerevisiae that were transformed with the indicated vectors were spotted on YPD plates with and without 12.5µg/mL anisomycin. Vector-derived yeast Drg1 (Rbg1) expression restores cell viability to near-WT levels and cells expressing bacterial and archaeal Drg (ObgE and mDrg) exhibit higher cellular growth rates than cells supplemented with the native yeast Drg1.

Drg homologs from all species share the conserved G domain (G1-G5), where a conserved serine residue of the G1 motif (S79 in yeast Rbg1) was shown to be critical for GTP-hydrolysis activity (Francis et al., 2012). A related yeast Rbg1 mutant containing alanine substitutions (including at the S79 residue) in the G1 motif (G77A, K78A and S79A) diminishes the association of the Rbg1/Tma46 complex with translating ribosomes (Daugeron et al., 2011). It is important to note that the S79N mutant of the native yeast Rbg1 failed to restore the cell viability in this assay (**Fig 1**. line 4), demonstrating that the GTPase activity of the Drg is critical to its cellular function.

Of note, presence of a more ancient Drg protein, such as ObgE or aDrg, to the Δrbg1 yeast showed more robust cell growth compared to the effect of native yeast Rbg1 supplementation. As shown in Figure 1 (line 6), addition of archaeal aDrg clearly restored cellular viability in the presence of anisomycin. Furthermore, expression of E*. coli Drg (ObgE)* (**Fig 1**. Line 6) restored cellular viability to a greater degree than the native yeast Drg1 (Rbg1) supplementation. By contrast, addition of another structurally similar E. coli translation factor, EF-Tu, failed to compensate for the lost function of yeast Drg1 (**Fig 1**. Line 7).

It is understood that evolutionarily ancient enzymes often perform more promiscuous functions compared to their modern counterparts. Therefore, it is not surprising to see better rescue of the yeast cell growth when a bacterial Drg is supplemented. However, human Drg1 was unable to restore cellular viability to the same degree as Δrbg1 yeast supplemented with WT yeast Rbg1 (**Fig 1**. line 8), indicating an increased level of functional specialization of the Drg protein over the course of evolution. Taken together, data from these straightforward experiments demonstrate a distinct and conserved GTPase-dependent cellular function shared among the Drg proteins from bacteria to eukaryotes.

### Translational functions of Drg proteins are conserved from bacteria to humans

Is the conserved cellular Drg function evident from the anisomycin growth assay related to cytoplasmic protein biosynthesis? We previously showed that yeast Drg proteins promote efficient translation when ribosomes slow down or pause on mRNAs (Zeng et al., 2021). To see whether Drg proteins share the same translational function in human cells, and to assess whether the bacterial homologs can also promote translation through such stall-inducing sequences, we monitored the degree of translation of a stalling reporter in Δdrg1 human cells expressing Drg1 orthologs from different species.

A dual-fluorescent reporter system, where EGFP (GFP) and mCherry (RFP) genes surround a poly(A) sequence (encoding poly-lysines, the K_20_ stalling sequence [**Fig. 2A**]), allows quantification of relative levels of translational stalling within cell populations at a single-cell resolution (Juszkiewicz and Hegde, 2017). According to the experimental design, ribosomes that stall at the poly(A) sequences are unable to proceed to translate the subsequent RFP gene, thereby decreasing the cellular concentration of RFP relative to GFP. As GFP is released prior to any ribosomal stalling event due to the 2A sequence, the amount of released GFP reflects the degree of translation on the reporter before translating ribosomes reach the stalling sequence. As such, the ratio of RFP/GFP fluorescence intensity obtained in each cell allows sensitive comparison of translation dynamics at such stalling sequences between cell populations. Translational machinery in Δdrg1 HEK293T cell populations exhibit a significantly decreased ability to translate past the poly(A) sequence compared to the WT cells expressing chromosomal Drg1, as indicated by a statistically significant ∼20% decrease in the average measured RFP/GFP ratio of Δdrg1 cells versus WT (**Fig. 2C**). This result suggests that human Drg1, similarly to its yeast homolog, Rbg1, promotes translation when ribosomes stall.

**Fig 2.**
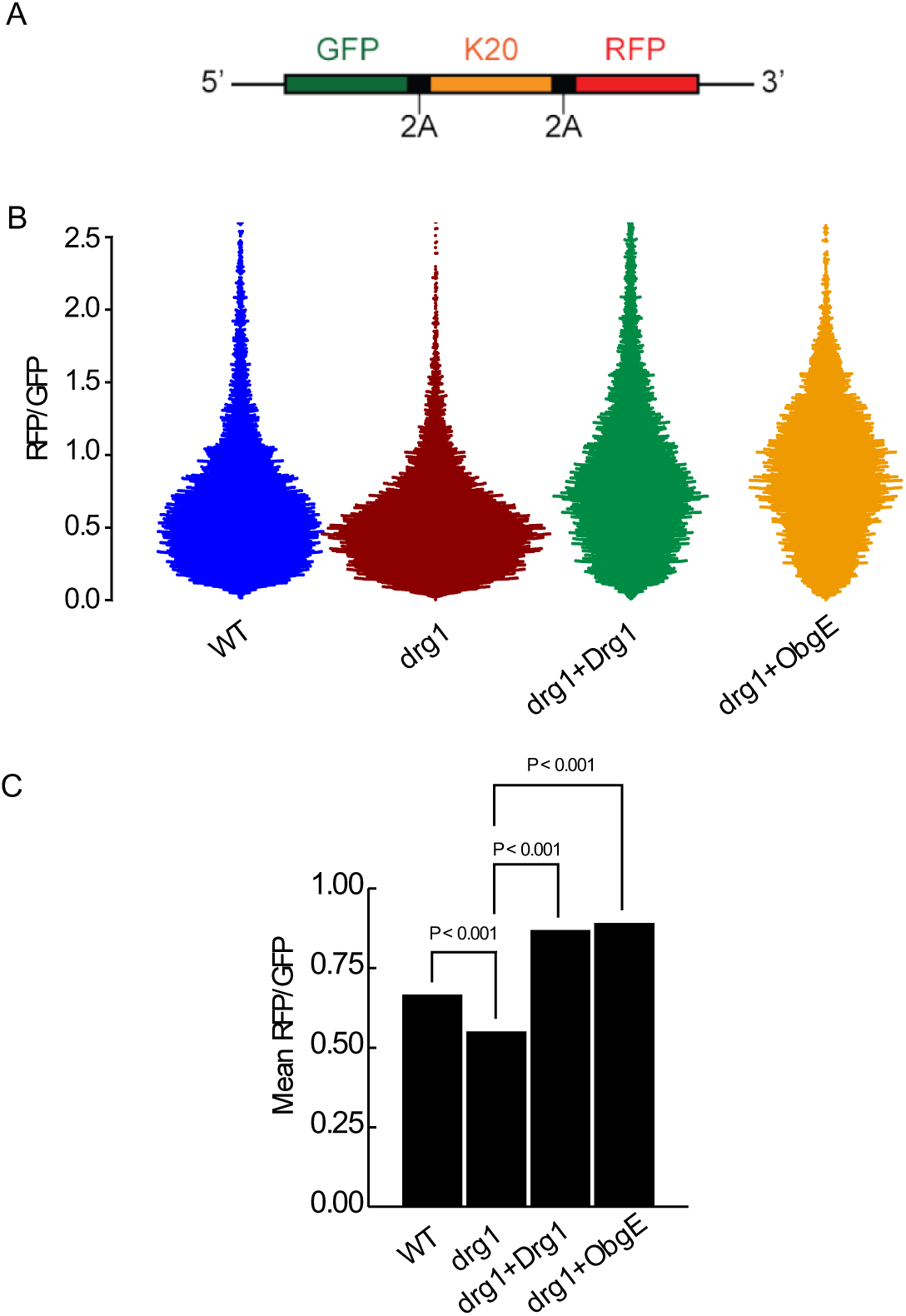
Function of Drg GTPases to promote translation in stalled ribosomes is conserved from bacteria to humans. **A. A schematic of the dual-fluorescent reporter used to monitor ribosomal stalling in human cells.** Dual-fluorescent reporter stalling construct encoding GFP (EGFP) and RFP (mCherry) flanking a polylysine (K_20_) stalling sequence. 2A sequences between each fluorescent protein and the stalling sequence ensure the release and accurate measurement of RFP and GFP protein to quantify ribosomal stalling at the K20 sequence via flow cytometry. **B. Bacterial ObgE restores the lost function of human Drg1 in promoting translation over the stalling sequence.** Beeswarm plot of RFP/GFP ratios illustrates that HEK293T cell populations expressing endogenous or exogenous Drg factors stall less frequently than Δdrg1 HEK293T cells. All WT and Δdrg1 HEK293T cells shown were transfected with a plasmid encoding the reporter construct. Cells were co-transfected with the reporter construct and Drg1 or ObgE overexpression vectors when indicated. Each datapoint represents the RFP/GFP ratio value from an individual cell. Approximately 8500 cells are shown for each condition reported here. C. Bar graph showing that translation is enhanced by the Drg GTPase. The mean RFP/GFP ratio from the indicated cell populations was shown. Δdrg1 HEK293T cell populations stall at the polylysine track significantly more than WT cell populations. Δdrg1 cell populations supplemented with hDrg1 or bacterial ObgE protein exhibit significantly more translation past the stalling sequence than Δdrg1 cell populations.

While chromosomal knockout of human Drg1 resulted in significant ribosomal stalling on the poly(A) sequence, translation through the stalling sequence was significantly restored when Δdrg1 cells were transfected with hDrg1- and ObgE-encoding vectors. As shown in Figure 2C, the average RFP/GFP ratios of Δdrg1 HEK293T cells increased by 58% and 62% upon expression of hDrg1 and ObgE, respectively. This observation shows that the function of Drg to promote efficient translation on stalled ribosomes is conserved across multiple domains of life. Moreover, overexpression of human hDrg1 and bacterial ObgE on a strong CMV promoter (See Methods) increased the average RFP/GFP ratios by roughly 30% compared to the average RFP/GFP ratio of the WT cells, indicating that cellular concentrations of these Drg GTPases influence ribosome dynamics in a dose-dependent manner.

### Drg proteins promote puromycin incorporation in stalled ribosomes

It is striking to see that bacterial ObgE supplementation alleviates human ribosomes stalled on the polyA sequence. In particular, eukaryotic Drg proteins bind to their partner protein and function as a heterodimeric GTPase on translating ribosomes (Francis et al., 2012; Zeng et al., 2021), so the robust functional replacement of eukaryotic Drg1 by monomeric bacterial ObgE is unexpected. Obg is an essential protein that plays diverse critical functions in bacteria (Kint et al., 2014; Leipe et al., 2002), such as its well-characterized ribosome biogenesis functions at the late stage of 50S biogenesis, where it oversees maturation of the GAC (Nikolay et al., 2021).

How do Drg proteins facilitate efficient translation on stalled ribosomes? Can Drg proteins bind directly to a stalled ribosome and promote it to continue protein synthesis past the stall? Biochemical and structural evidence obtained from the characterization of 80S ribosomes bound with yeast Drg1/Dfrp1 (Rbg1/Tma46) complexes led to the hypothesis that Drg proteins promote peptide bond formation by stabilizing a paused or stalled ribosome in a translationally productive conformation. To test this hypothesis, and to identify the possible mechanism of action of Drg GTPases in the ribosome, we studied puromycin reactivity in ribosomes that are stalled on a poly(A) sequence using a defined E coli *in vitro* translation system devoid of ribosome biogenesis, quality control, protein release, and recycling factors.

Since premature 3’ truncation of mRNA is a long-established means to stall elongating ribosomes reliably and efficiently at defined loci (Perara et al., 1986), we used an mRNA construct lacking an in-frame stop codon but containing 0 or 20 consecutive AAA codons at the 3’ terminus to stall elongating ribosomes with polylysine tracks occupying their exit tunnels (**Fig 3A**). This way, the translating ribosomes will stall at the end of the K0 mRNA due to lack of a stop-codon, cognate protein release factor, and no-go decay factors, or stall at the polyA sequence in the K20 mRNA.

**Fig. 3.**
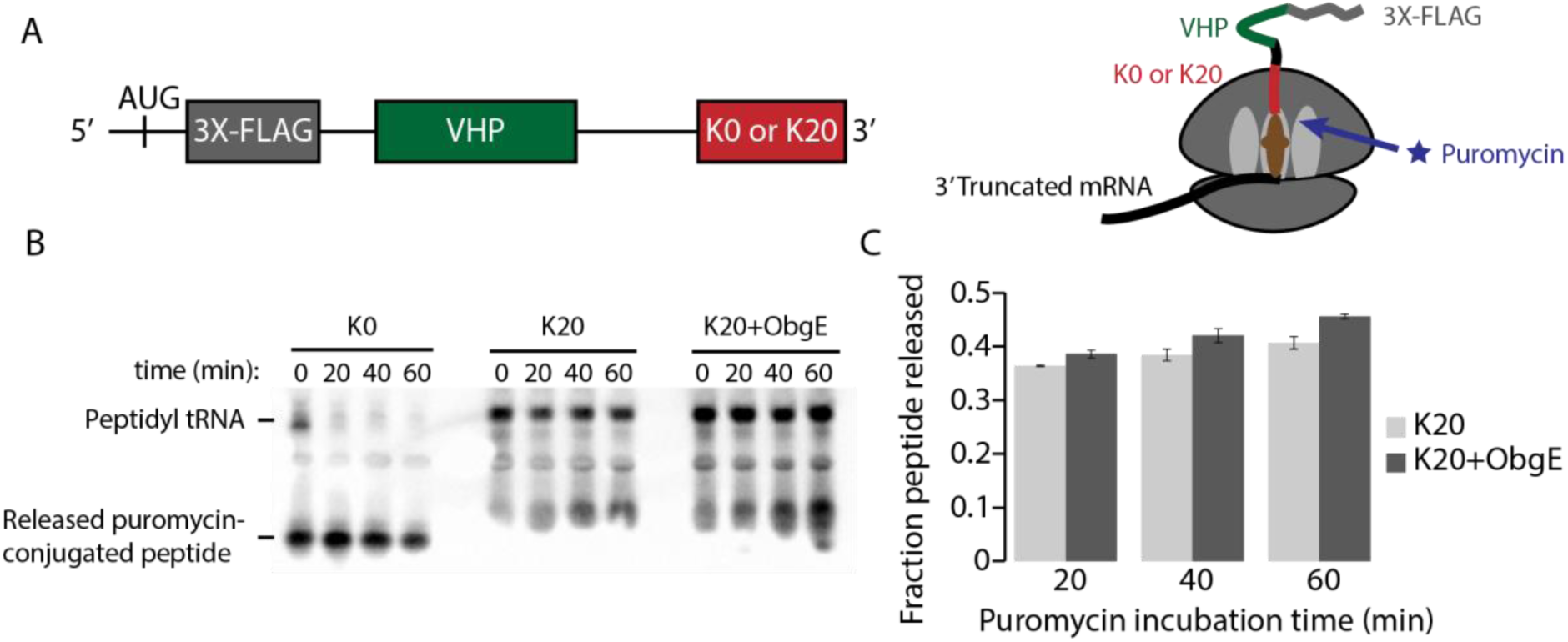
E.coli Drg, ObgE, facilitates peptide bond formation in stalled bacterial ribosomes A. Schematic of the mRNA construct used for the puromycin incorporation assay. Ribosomes were stalled at the truncated 3’ terminus of mRNAs containing either 0 (K0) or 20 consecutive (AAA) codons (K20) in a defined reconstituted *in vitro* E.coli translation system devoid of release, recycling, and quality control factors. 1mM puromycin was added to all reactions to measure peptidyl transferase function by quantifying puromycin incorporation which leads to the subsequent release of puromycin-conjugated nascent peptides. 2μM recombinant ObgE or an equal volume of buffer were added concurrently with puromycin to assay the ability of ObgE to facilitate peptide bond formation in the ribosomes stalled while translating the test sequences. **B and C. Increased puromycin incorporation in the K20-stalled ribosome in the presence of ObgE and GTP.** Representative western blot. The amount of puromycin incorporated into the nascent peptides was visualized and quantified by αFLAG immunoblotting (B). Quantification of puromycin incorporation and release from three replicates of the *in vitro* puromycin incorporation reactions of ribosomes stalled while translating the K20 sequence. Increased peptidyl transferase activity is observed upon incubation with recombinant ObgE at 20, 40, and 60 minutes after the addition of puromycin. Error bars for each group indicate the standard error of the mean (C).

It was shown that positively charged lysine stretches encoded by the polyA sequence induce conformational changes in the PTC that disfavor peptidyl transfer (Chandrasekaran et al., 2019; Tesina et al., 2020). To assess whether Drg proteins alleviate translational stalling by promoting elongation-competent conformational rearrangements in the ribosome, we monitored puromycin incorporation by stalled ribosomes. As a transition state analog, puromycin has long been used as a direct measure of the geometric optimality of the PTC. It mimics the CCA-end of an aminoacylated tRNA and can bind to the ribosomal A site. The reactive moiety of puromycin contains the requisite nucleophilic amine to attack the carbonyl carbon adjacent to the ester bond connecting the nascent peptide to the tRNA in the ribosomal P site in a manner identical to peptidyl transfer. Therefore, comparison of the levels of puromycin incorporation into nascent chain complexes within different ribosomal complexes will inform about relative differences in the capacities of their active sites (i.e., the PTC) to catalyze peptidyl transfer. For example, if the nascent chain conformation disfavors peptide bond formation due to a positively charged polylysine track in the exit tunnel, reduced puromycin reactivity is observed (Chandrasekaran et al., 2019). In the presence of Drg proteins, increased puromycin reactivity will demonstrate favorable conformational changes in the PTC that are induced by this GTPase to compensate for the suboptimal substrate orientation within a stalled ribosome.

Using an E. coli-derived *in vitro* translation system that contains *only* initiation and elongation factors, we assayed whether ObgE can target stalled ribosomes to promote puromycin incorporation. As expected, ObgE demonstrated a consistent ability to facilitate puromycin incorporation in ribosomes that were stalled while translating the poly(A) sequence. As shown in Fig 3, comparing the translation of 3’truncated mRNAs lacking poly(A) encoded polylysine tracks (the K0 control) with mRNAs encoding 20 contiguous lysines (K20) (**Fig. 3B**), the ability of K20- stalled ribosomes to catalyze puromycin incorporation was severely impaired. Importantly, ribosomes translating the K20 sequence in the reaction mixtures containing equimolar ObgE:70S concentrations exhibited a 6-12% increase in puromycin incorporation over the time course of 20-60 min when compared to otherwise identical reactions lacking ObgE (**Fig. 3C**). Since the amount of puromycin incorporation serves as a direct indicator of the level of peptide bond formation in the ribosome, this result demonstrates that the bacterial Drg ortholog, Obg, facilitates peptidyl transfer in the stalled ribosomes.

## Discussions

In this study, we show the essential bacterial GTPase Obg is an ortholog of eukaryotic Drg proteins, and the GTP-dependent activity of this protein to alleviate stalled translation is conserved from bacteria to humans. How does the conserved GTPase activity in Drg proteins contribute to increased peptidyl transfer in stalled ribosomes? Based on our results and the current knowledge in the field, we propose a processive model for Drg function in translation (**Fig. 4**). In this model, Drg proteins help the “temporarily troubled” ribosomes to resume translation by playing a molecular “engine” role: the GTP hydrolysis in a Drg facilitates conformational rearrangement within the ribosome to promote peptide bond formation and the subsequent dissociation of the Drg from the ribosome. In doing so, the Drg proteins decrease the likelihood that the ribosome that they act on is collided with when its translation slows down, or decrease the time that the ribosome is in the “collided disome” state if it has already been collided with by the trailing ribosome. Both scenarios will alleviate translation stalls and actively remove the collided disome, which is the substrate of ZNF598 and downstream quality control pathways (Kim and Zaher, 2022; Meydan and Guydosh, 2021; Wu et al., 2020), thereby promoting the processivity of translation on mRNA.

**Fig 4.**
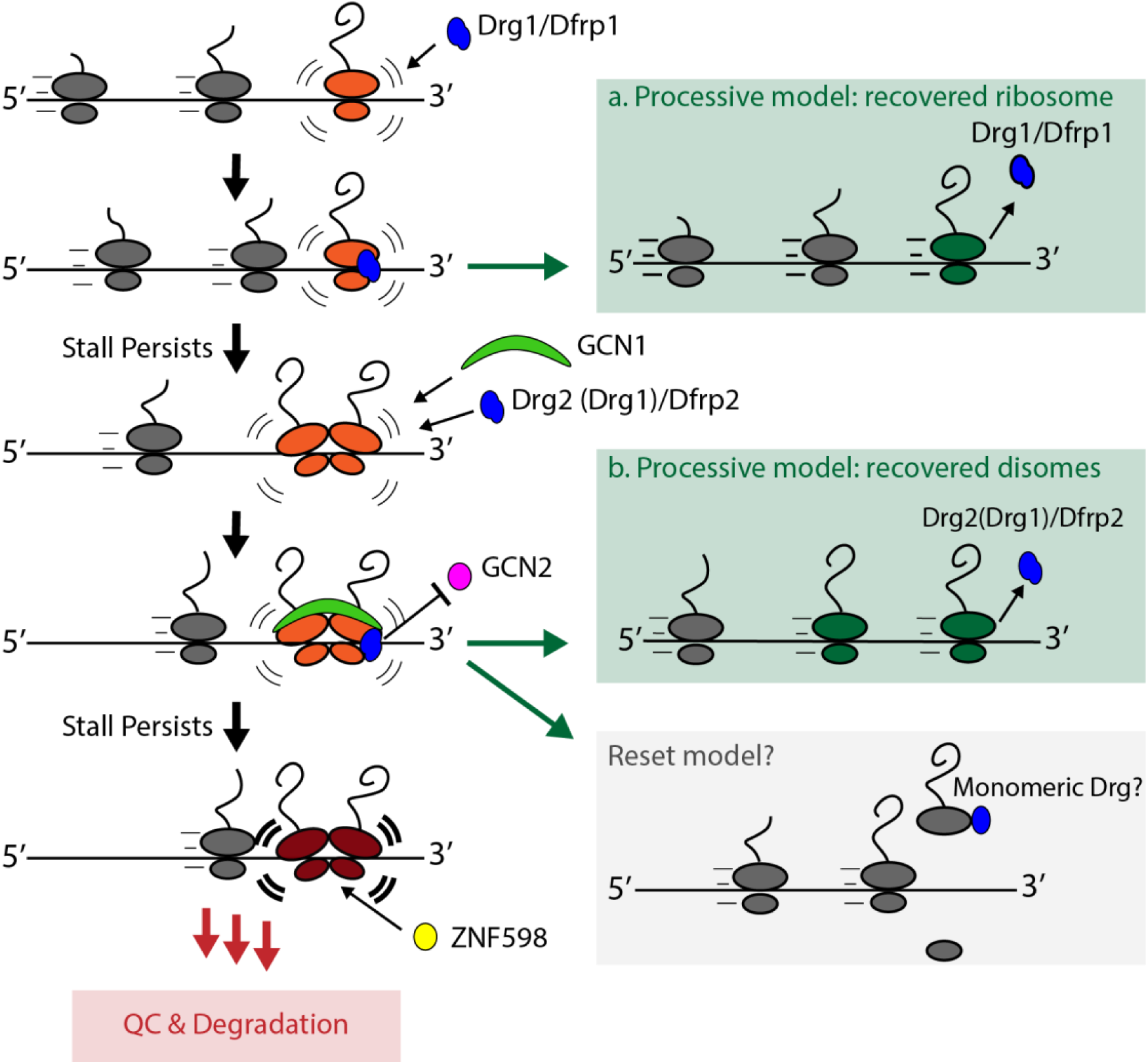
Proposed model for Drg function in translation. Here we propose two distinct roles for the two eukaryotic Drg proteins: Drg1 and Drg2. Both Drg1 and Drg2 bind to stalled ribosomes to promote the resumption of translation, rather than downstream quality control and stress response pathways, thereby providing the ribosome an opportunity to overcome the stall before mRNA translation is prematurely and unproductively terminated. Drg1 binds to slowed or stalled translating ribosomes which have not been collided with, promoting these ribosomes to resume translation of their mRNAs (A, processive model). If these ribosomes are stalled for a longer period, trailing ribosomes will collide with them. The formation of collided disomes are targets for known quality control and stress pathways such as NGD, RQC, ISR, and the ribotoxic stress response (RSR). Drg2/Dfrp2 target the collided disome by binding to GCN1, providing the problematic ribosome with more time to ‘decide’ whether to commit to quality control, all the while promoting the leading ribosome in the collided pair to continue translating (B, processive model). Of note, Drg proteins can attenuate the ISR pathway by competing with GCN2 for binding to the ribosome in a productive way (i.e., place-holder model). Finally, we cannot rule out the possibility that Drg proteins may, at some point in this pathway, target the problematic ribosomes for splitting in a manner that preserves the integrity of the involved mRNA (grey box, the reset model).

Several lines of evidence support the processive model. First and most importantly, we show that bacterial Drg stimulates puromycin incorporation within stalled ribosomes, providing direct evidence to support the translation-promoting activity of conserved Drg GTPases is indeed directly related to peptidyl transfer, the core catalytic function of the ribosome which is demonstrably impaired within poly(A)-stalled ribosomes. Second, the presence of yeast Drg1 (Rbg1) increases the stability of reporter mRNAs harboring a strong stalling sequence (Zeng et al., 2021), suggesting that the stimulation of translation by Drg factors diverts stalled ribosome complexes away from quality control pathways which necessarily degrade the affected mRNA. Third, a recent cryoEM structure of yeast Drg1/Dfrp1 bound to the 80S ribosome reveals that these ribosomes assumed entirely translation-competent conformations in their PTCs and decoding centers (Zeng et al., 2021). In contrast, structural analyses on polyA-stalled 80S ribosomes, as well as collided disomes formed on a synthetic mRNA with strong stalling sequences show that these “deeply troubled” ribosomes are functionally unable to resume translation and are thus committed to the ribosome-associated quality control pathway (Best et al., 2023; Chandrasekaran et al., 2019; Ikeuchi et al., 2019; Juszkiewicz et al., 2018; Matsuo et al., 2020; Tesina et al., 2020). Fourth, results from our mass spectrometry data show that the pool of proteins associated with Drg-bound translating ribosomes are enriched with chaperones and other nascent chain-interacting proteins but not the known quality control factors.

Fifth, an earlier genetic study shows that one of the Drg complexes, Drg1 or Drg2, is required for optimal growth in yeast lacking the Slh1 gene (Daugeron et al., 2011), suggesting that Drg proteins and Slh1 are most likely to function in different cellular pathways that have related functions. The protein product of this gene, Slh1 protein, is an ATP-dependent helicase that dissociates the two subunits of a stalled leading ribosome in a collided disome to commit it to ribosome-associated quality control (Best et al., 2023; Ikeuchi et al., 2019; Juszkiewicz et al., 2020; Matsuo et al., 2017; Matsuo et al., 2020). Therefore, Drg proteins likely play a role that is mechanistically different, but related to ribosome-associated quality control.

It is important to note that in addition to the “engine” role, Drg proteins likely also play a related “place holder” role, where ribosomal recruitment of a Drg helps to prevent premature activation of stress responses and quality control. A recent structural study shows that yeast Drg2/Dfrp2 associates with collided disomes wherein Dfrp2 contacts Gcn1 protein that binds to the two collided ribosomes (Pochopien et al., 2021). In this interaction, the conserved RWD domain of Dfrp2 interacts with GCN1 in a way that is suggested to occlude GCN2 from binding GCN1 (Pochopien et al., 2021; Wout et al., 2009), an interaction that is necessary for activation of the integrated stress response (Garcia-Barrio et al., 2000).

It is worth noting that we cannot entirely exclude a reset model, where the Drg GTPase helps to disassemble the stalled ribosomes to clear the roadblock on the mRNA. E. coli ObgE was reported to bind to the 50S GAC and was suggested to serve as an anti-association factor, preventing premature association of the ribosomal small subunit before completion of GAC maturation in pre-50S subunits (Feng et al., 2014; Nikolay et al., 2021). However, it is important to note that the two models, the processive and reset models, need not be mutually exclusive. Furthermore, we have shown that yeast Drg1 binds to tRNAs in the cell (Zeng et al., 2021). Since there are other processes, such as tRNA selection, proofreading, and accommodation, occurring in the ribosome prior to the peptidyl transfer reaction leading to the peptide bond formation, these steps can also be facilitated by the Drg GTPases, especially tRNA accommodation, which is often rate-limiting. Taken together, our data demonstrate that Drg proteins are a new general translation factor that directly affords cells from three domains of life a new form of translation control. Detailed functional and mechanistic investigations into the Drg GTPases of eukaryotic, archaeal, and bacterial species represent exciting areas for future study.

## Acknowledgments

We thank Dr. V. Ramakrishnan for sharing the original mRNA construct used in the puromycin incorporation assay in mammalian cells, and Dr. Vish Chandrasekaran from the Ramakrishnan laboratory for insightful discussions. We thank Dr. Gary Olsen for helping us with an extensive phylogenetic analysis of the Drg proteins from bacteria to humans. We thank Dr. Toshifumi Inada for a helpful discussion on the possible functional overlap between Rbg and Slh1. We also thank members of the Jin Lab, including Sounak Saha, for helpful discussions and experimental help, and former lab member Dr. Melissa Pires-Alves for guidance and contributions to the project which have facilitated this publication. We thank the Roy J. Carver Biotechnology Center for access to the flow cytometry facility. This research was funded by the startup fund to H. Jin from the University of Illinois at Urbana-Champaign.

## Methods

### Cell lines, strains, and Growth Conditions

The BY4742 strain of *Saccharomyces cerevisiae* (*MATα his3Δ1 leu2Δ0 lys2Δ0 ura3Δ0)* was used in this study. Yeast chromosomal knockouts were previously generated using homologous recombination techniques (Baudin et al., 1993; Longtine et al., 1998; Zeng et al., 2021). Δrbg1 BY4742 S. cerevisiae used in this study have the following genotype: *MATα his3Δ1 leu2Δ0 lys2Δ0 ura3Δ0 rbg1::Kana*, and were grown in YPD+G418 medium unless indicated otherwise. When indicated, S. cerevisiae were transformed with recombinant vectors via the established LiAc/ssDNA/PEG method, with minor modifications (Gietz, 2014).

Adherent HEK 293T cells used in this study were cultured in DMEM medium supplemented with 10% FBS, 1mM Pyruvate, and 100U/mL Penicillin + 100μg/mL Streptomycin and grown in a 37° incubator containing 5% CO_2_. HEK 293T Drg1 genomic knockouts were generated by co-transfection of vectors encoding Cas9 and gRNAs to target the indicated genes.

### Protein Expression and Purification

BL21(DE3) cells were transformed by a standard heat shock method with a pET28-a vector expressing WT ObgE (derived from BL21(DE3) E.coli) with an N-terminal 6X-His tag. LB medium was inoculated with overnight cultures and was grown at 37° with 220RPM shaking until it reached an OD_600_ value of 0.6-0.7. Expression was induced by addition of 300μM IPTG and cultures were incubated at 22° for 16hrs with 145RPM shaking. Cells were harvested by centrifugation, washed with 1XPBS, then resuspended in Lysis Buffer (20mM Tris-HCl, pH7.5; 500mM NaCl; 1.5mM MgCl_2_; 50mM Imidazole; 1X cOmplete EDTA-free Protease Inhibitor (Roche); 40U DNaseI (NEB)). Cells were lysed via a French press, then lysates were clarified by centrifugation. Clarified lysates were incubated with Ni-NTA Agarose (Qiagen) at 4° for 1.5 hr, then applied to a gravity column and washed with Lysis Buffer lacking protease inhibitor and DNaseI. Protein was eluted in three successive applications of Elution Buffer (20mM Tris-HCl, pH7.5; 500mM NaCl; 1.5mM MgCl_2_; 500mM Imidazole), then concentrated with 30kDa MWCO Amicon centrifugal filters (Millipore Sigma). Concentrated fractions were further purified via a Superdex 200 10/300 Size-exclusion column in an Åkta FPLC (GE) in ObgE Storage Buffer (20mM Tris-HCl, pH7.5; 150mM KCl; 5mM MgCl_2_; 2mM 2-mercaptoethanol), concentrated, and flash frozen in liquid nitrogen and stored at −80°.

### Anisomycin Cell Viability Assay

A yeast spot assay using serial dilutions was used with modifications (Xu et al., 2014). WT and Δrbg1 BY4742 cells were transformed with plasmids encoding Drg proteins from different species and E.coli EF-Tu that were expressed under a PGK promoter. Transformed cells were grown to lag phase at 30° with shaking in SC-ura medium (MP Biomedicals) and WT cells were grown in YPD medium. Fresh YPD medium was inoculated with the cells to an OD_600_ value of 0.2, then cells were grown until mid-log phase (approximately OD_600_ 0.6) after which cells were harvested by centrifugation, then resuspended at an OD_600_ of 1.0 in 1X PBS. Serial dilutions were made as indicated, then 1μL of each was spotted on YPD and YPD+12.5μg/mL Anisomycin plates, which were grown at 30° for the indicated times.

### In vivo Flow Cytometry Stalling Assay

Actively dividing WT and Δdrg1 HEK293T cells were trypsinized and passaged to cell culture-treated 6-well dishes (Thermo) at a cellular density of 2×10^5^ cells/mL. After 24hrs, cells were transfected with plasmids encoding the dual-flourescent reporter mRNA and, when indicated, plasmids encoding Drg proteins on strong CMV promoters using the Transit293 transfection system (Mirus Bio) according to the manufacturer’s recommendations. Cells were harvested 24hr after transfection, then were washed twice with and stored in ice-cold 1X PBS. A BD LSR Fortessa flow cytometer was used to selectively excite and measure emission for both cellular RFP (mCherry) and GFP (EGFP). FCS Express 7 was used to gate and isolate events from individual cells, then the beeswarm plot was generated with the “Beeswarm” R package. P-values were calculated for the comparison of means between cell populations using the Mann Whitney U test.

### Puromycin incorporation Assay

E. coli in vitro translation was used to measure puromycin reactivity in stalled ribosomes. The experiment was conducted similarly to a previously published method using mammalian translation systems (Chandrasekaran et al., 2019) with modifications. The vector encoding the 3’-truncated mRNA used to assay puromycin incorporation in ribosomes harboring polylysine tracks was modified from a vector generously gifted by the Ramakrishnan laboratory of the MRC LMB in Cambridge. T7 promoter and Shine-Delgarno sequences were added at the 5’ end, and the 3’ end was truncated and modified to encode zero lysines (for the K0 construct) or twenty consecutive lysines by PCR (for the K20 construct). In addition to the 3’ test sequences, the recombinant DNA encodes an N-terminal 3XFLAG tag, and a central villin headpiece domain. The mRNA construct was co-transcriptionally translated using the defined, reconstituted E.coli PURExpress ΔRF123 expression system (NEB) at 37° for 1hr. Expression was halted by placing the reactions on ice, then 2 μM ObgE or an equal volume of ObgE Storage Buffer and 1mM puromycin were added. Puromycin-treated translation reactions were returned to 37 °C. Aliquots were taken at the indicated time points and quenched by diluting in SDS-PAGE loading buffer and incubating on ice. Puromycin-reacted and released peptides were separated from peptidyl tRNAs by SDS-PAGE and then were visualized via anti-FLAG immunoblotting. Percentages of released puromycin-conjugated peptides were quantified for each replicate using ImageJ (NIH) integration of corresponding band intensities.

### SDS-PAGE and Immunoblotting

Protein samples were analyzed by standard SDS-PAGE and immunoblotting techniques. For routine SDS-PAGE, proteins were electrophoresed through homemade 12% polyacrylamide (Biorad) gels running at a constant 200-220V, and stained with Coomassie blue. For immunoblotting, NuPAGE 4-12% Bis-Tris gels (Thermo Fisher) were loaded with protein samples and SDS-PAGE was conducted at a constant 180V for approximately 1hr. Protein samples were transferred to nitrocellulose membranes (Amersham) upon which M2 Anti-FLAG mouse antibodies (Sigma) were used to visualize 3X-FLAG fusion peptides.

